# *Connectome*: computation and visualization of cell-cell signaling topologies in single-cell systems data

**DOI:** 10.1101/2021.01.21.427529

**Authors:** Micha Sam Brickman Raredon, Junchen Yang, James Garritano, Meng Wang, Dan Kushnir, Jonas Christian Schupp, Taylor S. Adams, Allison M. Greaney, Katherine L. Leiby, Naftali Kaminski, Yuval Kluger, Andre Levchenko, Laura E. Niklason

## Abstract

Single-cell RNA-sequencing data can revolutionize our understanding of the patterns of cell-cell and ligand-receptor connectivity that influence the function of tissues and organs. However, the quantification and visualization of these patterns are major computational and epistemological challenges. Here, we present *Connectome*, a software package for R which facilitates rapid calculation, and interactive exploration, of cell-cell signaling network topologies contained in single-cell RNA-sequencing data. *Connectome* can be used with any reference set of known ligand-receptor mechanisms. It has built-in functionality to facilitate differential and comparative connectomics, in which complete mechanistic networks are quantitatively compared between systems. *Connectome* includes computational and graphical tools designed to analyze and explore cell-cell connectivity patterns across disparate single-cell datasets. We present approaches to quantify these topologies and discuss some of the biologic theory leading to their design.

## Introduction

Cell-to-cell communication is a major driver of cell differentiation and physiological function determining tissue/organ development, homeostasis, and response to injury. Within tissues, cells have local neighbors with whom they directly communicate via paracrine signaling and direct cell-cell contact, and long-range or mobile partners with whom they exchange information via endocrine signaling. In solid tissues, cell types have specific cellular niches incorporating localized matrix and signaling environments, which facilitate phenotypic maintenance and support specialized cell functions. Circulating immune cells use an extensive library of chemokines to coordinate multicellular system responses to threat or injury. The advent of single-cell technologies has made it technologically possible, for the first time, to codify cell-specific ligand-receptor patterns in complex tissues with both high accuracy and robust statistical confidence. The resulting signaling topologies are strongly suggestive of intercellular communication vectors and can be revealed through a variety of techniques (*1–8*). The combination of single-cell sequencing data with ligand-receptor mapping is therefore a promising approach to exploring, understanding, and reverse-engineering complex tissue systems-biology for biologic, therapeutic, and regenerative efforts.

A connectomic network from a single tissue has unique properties that must be taken into consideration for biologically relevant downstream information processing. Tissue-derived connectomic networks are directional – i.e. each ligand-receptor interaction matrix is asymmetric; multi-modal – i.e. many ligand-receptor mechanisms contribute to the connectome; and weighted – i.e. interaction edges can be assigned quantitative values. These properties make data mining and data visualization substantially more complex than in some other genres of network science. Added dimensions can additionally come into play when it is necessary to compare cell-cell signaling in tissues between experimental conditions, over time during growth or remodeling, or between disparate tissue systems in which the same cell type annotations are not necessarily present. Here we describe a new computational package in R called *Connectome* which facilitates each of these tasks.

*Connectome* is a multi-purpose tool designed to create ligand-receptor mappings in single-cell data, to identify non-random patterns representing signal, and to provide biologically-informative visualizations of these patterns. *Connectome* formalizes and generalizes the methods first developed in Raredon et al. 2019 (*1*), allowing extension of the same analyses to any single-cell dataset, or sets thereof, in association with the R package *Seurat*. By default, *Connectome* uses the FANTOM5 database of ligand-receptor interactions (*9*), but it also allows mapping against any user-provided ligand-receptor list. Because the reference database can be customized, *Connectome* can be used to investigate newly discovered, or hypothesized, ligand-receptor mechanisms of particular interest.

To demonstrate *Connectome*, we apply the software analysis in three distinct use cases. First, we demonstrate application to a single tissue by analyzing single-cell human pancreas data. Second, we describe differential connectomics, comparing IFN-stimulated human PBMCs with unstimulated control data. Third, we apply *Connectome* to a longitudinal wound-healing dataset in mouse muscle. For brevity, extended discussion of certain topics has been compiled in a Supplemental Methods sections. All software is publicly available at https://github.com/msraredon/Connectome. Detailed vignettes and instructions for use are published at https://msraredon.github.io/Connectome/. Exact scripts for replicating all figures in this manuscript are included in Supplemental Data D1.

### Defining the data structure of tissue connectomics

The connectomic mapping discussed here treats every cell parcellation (e.g., different cell types, phenotypically distinct cell states of the same cell type, etc.) as a single node, averaging ligand and receptor values across a given cell parcellation to yield mean values which are then linked. Mean-wise connectomics has the advantage of accommodating the zero-values intrinsic to single-cell data, while simplifying the system so that every cell parcellation is represented by a single, canonical node. However, as this approach will blend the effects of all cellular archetypes within a cluster, initial cell parcellation must be done carefully for the resulting connectomic networks to be biologically meaningful.

An edgeweight must be defined for each edge in the celltype-celltype connectomic dataset. *Connectome*, by default, calculates two distinct edgeweights, each of which captures biologically relevant information. The first edgeweight, also referred to as ‘w_1_’ or ‘weight_norm_,’ is defined as a function (by default, the product) of the celltype-wise normalized expression for the ligand and the receptor, or

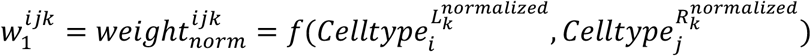

Where (k) denotes a specific ligand-receptor pair, 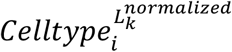 is the normalized expression of ligand *L*_*k*_ in celltype (i), and 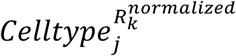 is the normalized expression of receptor *R*_*k*_ in celltype (j). Weight_norm_ is useful for differential connectomics, in which exact edges are compared across tissue conditions. However, Weight_norm_ is encumbered by the fact that it ignores the cell-type specificity of many ligands and receptors. Weight_norm_ will weigh a highly expressed but non-specific cell-cell link as stronger than a highly specific but lowly-expressed gene. This does not align with biologic intuition for many cell-cell signaling mechanisms, nor with the goal of parsing out rare cell types that may have outsized biological relevance. Therefore, *Connectome* also, by default, calculates a second edgeweight, alternatively referred to as ‘w_2_’ or ‘weight_scale_,’ which is defined as a function (by default, the mean) of the gene-wise z-scores of the ligand and receptor, or

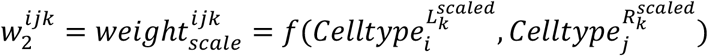

where 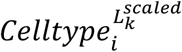 is the system-scaled expression of ligand *L*_*k*_ in a celltype (i) and 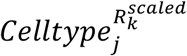 is the system-scaled expression of receptor *R*_*k*_ in a celltype (j). Because weight_scale_ incorporates information on the topology of the entire system being analyzed, leveraging system z-scores of each gene, it is ideally suited to exploring cellular ‘roles’ within a single tissue system. It also can be used to compare cellular prominence within signaling networks across disparate tissue systems, or across species, in which normalized values may vary but their *relative* expressions across cell types do not (*1*).

Alternative edgeweights, such as those incorporating a score for likelihood of downstream signal transduction (*5*) or those which take into account proximity information from spatial transcriptomics or histology-registered techniques (*10, 11*), may reveal additional patterns. For the purposes of this software program, *Connectome* defaults to the above edgeweights definitions because we have found them to be effective for understanding biological questions addressed in our studies.

Statistical significance depends upon the question being asked, the test being applied, and the threshold for important differences, and has a very different meaning depending on whether a research team is investigating a single tissue, comparing multiple tissues, or studying a physiologic process. However, we find two general statistical patterns to be of key importance, and we have built-in functionality to *Connectome* to help the researcher narrow in on these patterns. First, when studying a single tissue system (i.e., one organ in one condition), it is often desirable to focus on those ligand-receptor interactions which are highly associated with (markers for) specific celltype-celltype vectors. While such a pattern is not a guarantee of biological relevance, it is often useful to learn those mechanisms which can mark celltype-celltype vectors in the same ways that transcripts can mark celltypes. Second, when comparing two tissue systems (i.e., one organ in two experimental conditions) it is generally interesting to focus on those mechanisms which, regardless of their association with a specific celltype-celltype vector, are differentially regulated across condition. In the online vignettes, we detail computational methods which can be used to define statistically significant edges matching each of these use cases.

### Visualizing the connectomic signature of a single tissue

Figure 1 shows the workflow for ligand-receptor connectomics in a single tissue. In this instance we use single-cell data from 8 separate single-cell RNA libraries of human pancreatic tissue, available through the *SeuratData* database (*12*). As a first step (Figure 1A), the tissue data are parcellated into defined cell types using a standard clustering workflow. Then, a mapping is created against a ground-truth database of known ligand-receptor mechanisms. This mapping yields a large edgelist, wherein nodes are defined as cell types and edge attributes contain quantitative information derived from cell type-specific expression levels. Source nodes denote the ligands, and target nodes refer to the cognate receptors. This data architecture is stored as a data frame in R, the environment which is particularly amenable to subsetting networks of interest and working with graph theory-based computational packages. This process is performed by a single function in *Connectome* which allows customization of edgeweight definition and edge attribute calculations.

**Figure 1:**
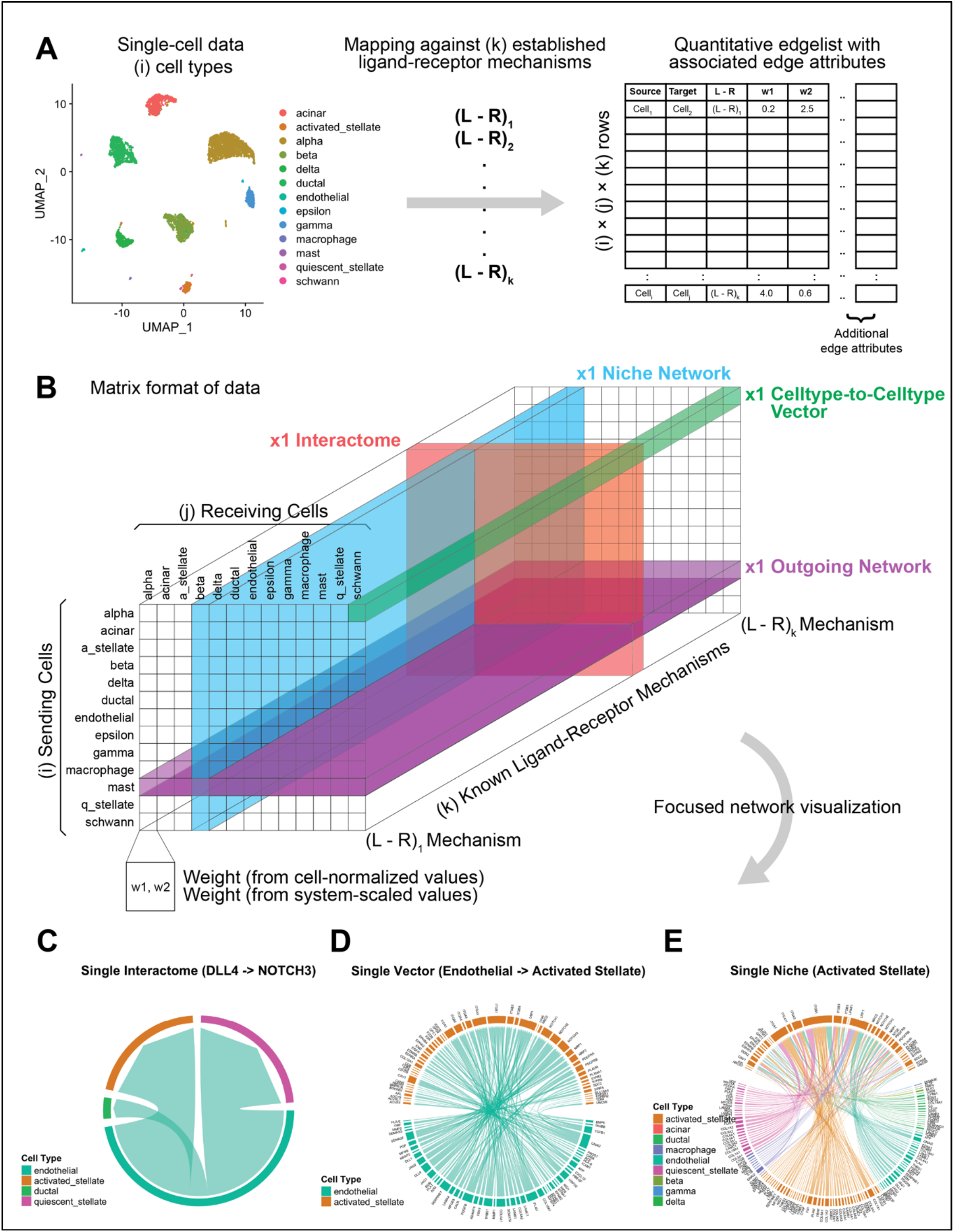
Single-tissue connectomic topology. (A) Pancreas single-cell data is parcellated into defined cell types. These cell types are then mapped against a ground-truth database of known ligand-receptor mechanisms. This yields a comprehensive edgelist of cells expressing ligand connecting to cells expressing receptor, with associated edge attributes. Note that a single ligand may hit multiple receptors and vice-versa (B) A conceptual visualization of the data architecture and biologically informative cuts through the data. (C-E) Selected quantitative visualizations of interactome-, vector-, and niche-networks made with *Connectome.* These three plots, respectively, allow (C) identification of top cell types utilizing a given L-R mechanism, (D) top mechanisms employed by a cell-cell vector, and (E) top cell-mechanism combinations in a position to influence a receiving cell. In C-E, edge thickness is proportional to weight_scale_, which is larger when an edge is more highly associated with a specific celltype-celltype vector. In all cases, the network shown has been limited to those edges in which the ligand and receptor are both expressed in > 10% of their respective clusters and have a p-value of < 0.05. In subpanel E, for illustration, the network has been further thresholded to those edges with a ligand and receptor z-score of > 1.

Conceptually, the data architecture for a single edge attribute (one column in the above discussed edgelist data frame in Figure 1A) can be thought of as a 3D matrix (Figure 1B), where rows are source (sending) cells, columns are target (receiving) cells, and the z-axis is the full list of ground-truth known ligand-receptor mechanisms. This allows clear visualization of the data that needs to be subset in order to explore a single interactome (red), outgoing network (purple), niche network (blue), or cell-cell vector (green). The blue rectangle represents a niche-network, containing information on all edges in a position to influence a single cell type. The purple rectangle shows all edges coming from a single cell type. The red rectangle is a single interactome, containing information for a single signaling modality between all cell types. The green prism represents a single cell-to-cell vector, containing information for all signaling modalities.

Visualization of these connectomic networks can be done in multiple ways. *Connectome* includes a series of functions designed for tissue network exploration, including one which generates plots allowing immediate quantitative visualization of individual interactomes (e.g. Fig 1C), celltype-to-celltype vectors (Fig 1D) and niche networks (Fig 1E). Further, the similarity between individual celltype-to-celltype vectors (vectortypes) can be analyzed using a k-nearest-neighbor style embedding (Fig S1). This style of visualization places vectortypes in a 2-dimensional space and provides a quantitative way to cluster vectortypes based on which ligand-receptor mechanisms are most highly weighted in each celltype-to-celltype pairing. We provide a custom function in *Connectome* to perform this analysis.

### Tissue network centrality analysis

A major goal of tissue science and cell biology is to understand the roles that individual cells types play and their ability to affect other cell types. It is of interest, therefore, to rank cell types based on their ability to produce or receive information within a given signaling network, or signaling family. To quantify these roles, *Connectome* is capable of performing a centrality analysis, an example which is shown in Figure 2. In this analysis, the total connectome for a single tissue is first subset down to only those edges belonging to a single signaling family (Figure 2A). This weighted graph is then used to calculate two centrality metrics: the Kleinberg hub and Kleinberg authority scores (represented as the dot size), and the cumulative directed edgeweight (x-axis), for each cell type within the network. The dot sizes arising from the Kleinberg scores are then used to visualize outgoing and incoming centrality of each cell type (Figure 2B), for a variety of specific signaling mechanisms. For example, the endothelial cell population of the pancreas has a high outgoing centrality score for signaling along the PDGB axis to the mesenchyme compartment (green circle in upper left panel of Figure 2B, “endothelial”, highly connected to the ochre circle in upper right panel, ‘activated stellate’). The result is a ranking of outgoing and incoming centrality for every cell type, across every signaling family, for each target cell type within the tissue. Collectively, this analysis creates a comprehensive portrait of potential extracellular signal transfer within a given tissue system. As shown in Figure 2, this information can then be used to group each signaling family based on the cell type that is best positioned to receive information (shown, sorted by incoming centrality) or best positioned to generate a signal (not shown - requires sorting by outgoing centrality).

**Figure 2:**
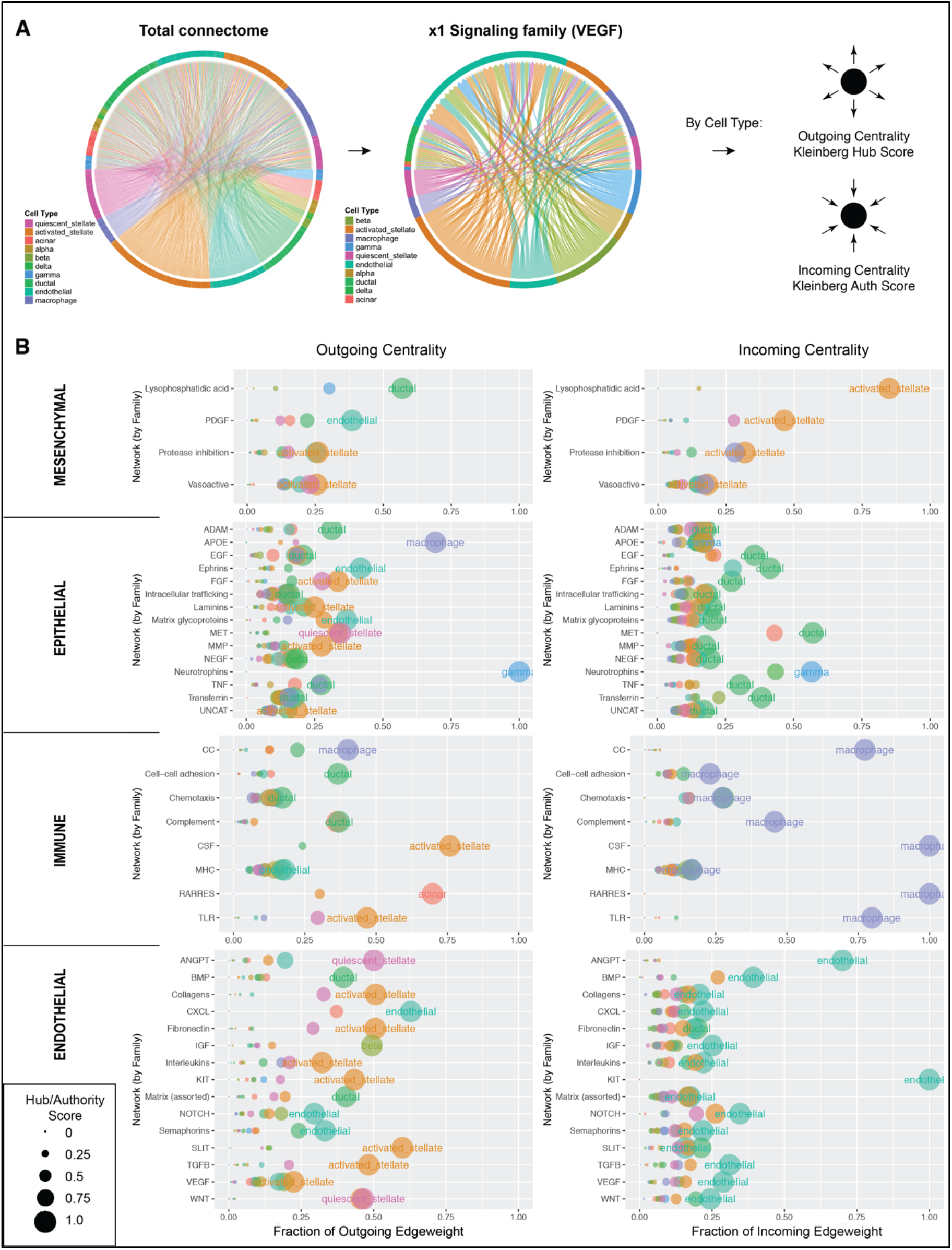
Centrality analysis for a single tissue system. (A) The global connectome for the pancreas is iteratively subset to each individual signaling family, after which Kleinberg hub and authority scores are calculated for each cell type. (B) Signaling families, grouped by dominant receiving nodes, crafts a quantitative portrait of global tissue system signaling architecture. Of note and in alignment with biologic intuition, mesenchymal cells are top network receivers of PDGF-family signals, epithelial cells are top network receivers of EGF-family signals, immune cells are top networks receivers of CSF- and CC (Chemokine)-family signals, and endothelial cells are in top network receiving positions for VEGF-, and ANGPT-family pathways. Centrality analysis was performed once for all signaling families, groupings were designated by top receiving node, and the four grouped plots were then individually re-made (see Supplemental Data D1).

### Differential connectomics with aligned nodes

It is common in systems biology to examine changes in cell-cell signaling between two systems, i.e., in a multicellular tissue before and after treatment with a drug or chemokine (Figure 3A). In one such example, if the same cell types are present in both systems, we may calculate direct, one-to-one comparison of all edges using the *Connectome* package. It should be noted, however, that within a differential connectome, there are *four* distinct types of perturbed edges, since for each cell in a differential edge, the ligand and the receptor may be either increased or decreased (Figure 3B).

**Figure 3:**
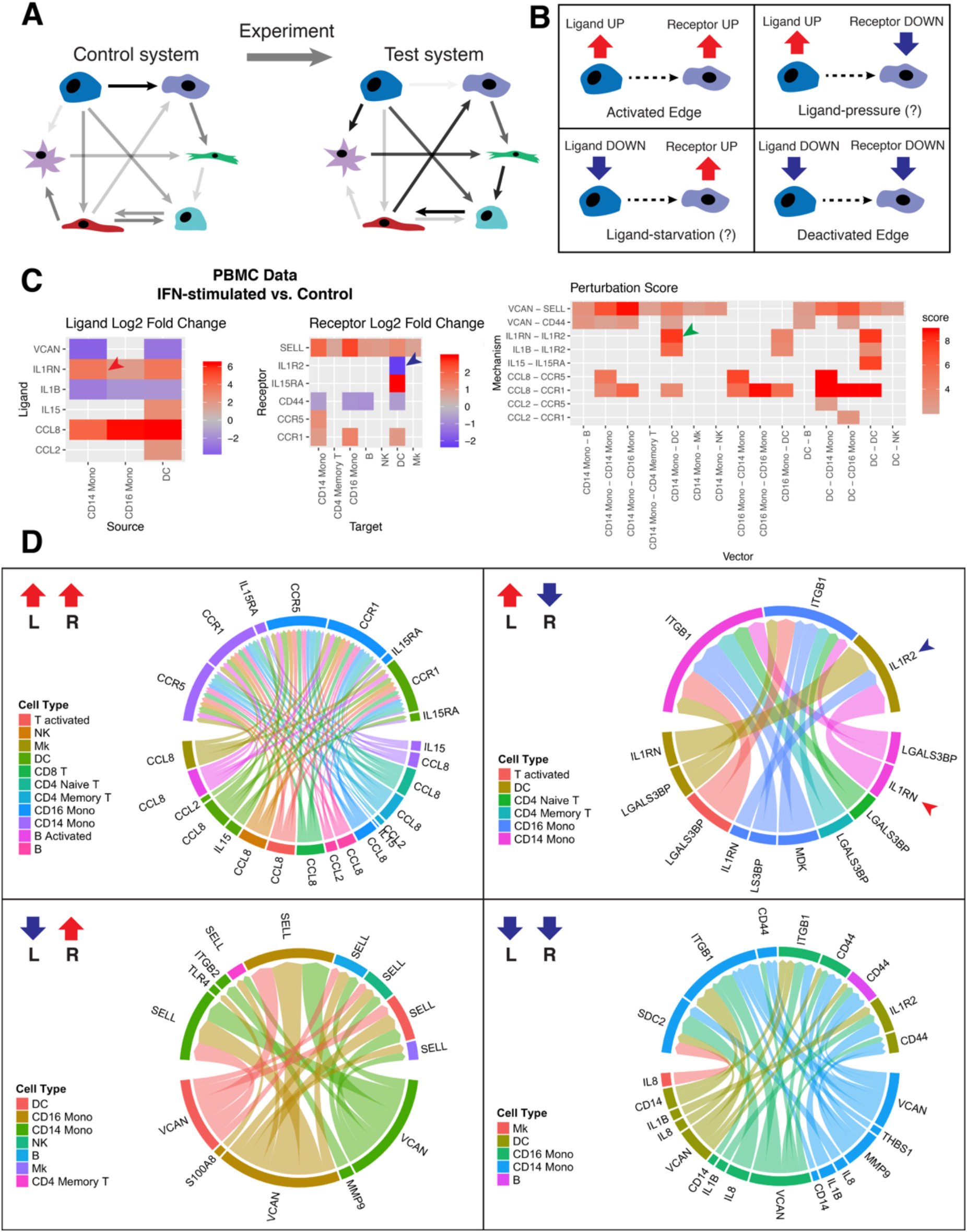
Differential tissue connectomics. (A) Schematic showing comparison of a perturbed multicellular system against a known control or reference set of interactions. (B) Assuming we are only interested in those edges in which either the ligand or the receptor change due to perturbation, each edge in a differential systems comparison falls into one of four distinct styles: the ligand and receptor are either both up, both down, or some combination. (If edges are also to be considered in which *only* the ligand or receptor change, then there are eight distinct categories of edge shift.) Dual ligand/receptor increase or decrease are consistent with edge activation or deactivation, respectively. A decreased ligand paired with an elevated receptor suggests ligand starvation, as is often seen in *in vitro* experiments. And increased ligand paired with a decreased receptor suggests the converse, ligand pressure. (C) Application to IFN-stimulated versus control PBMC data, showing how a positive ligand fold change (red arrow) and negative receptor fold change (blue arrow) combine to form a single positive edge perturbation score (green arrow). For illustration purposes, the differential network here has been heavily thresholded (minimum perturbation score of 2, a minimum expression cutoff of 20%, and only significant edges) yielding the presence of grey squares and the low number of displayed nodes and mechanisms. (D) Shows differential cell-cell signaling in IFN-stimulated PBMCs versus controls, sorted by style of perturbation. Edge thickness is proportional to perturbation score. Edge color correlates with source celltype. Blue and red arrow-heads emphasize the same edge similarly emphasized in (C). Networks in D, for illustration, have been thresholded to a minimum perturbation score of 2 and a minimum percentage cutoff of 10%. In all cases, differential network analysis has been limited to those edges where the expression of both the ligand and receptor, in their respective populations across condition, are differentially expressed with a p-value < 0.05 as assessed by a Wilcoxon Rank Sum test (see Methods, Supplemental Data D1).

If both the ligand and receptor are upregulated, we may reasonably call the edge ‘activated’, while conversely, when both the ligand and receptor are downregulated, we may think of the edge as ‘deactivated’. The two alternate cases, when a ligand is upregulated and a receptor down, or vice-versa, are more complicated to interpret. Although such patterns may initially suggest ligand-pressure or ligand-starvation, in which the downstream cell changes its character in response to upstream influence, it is important to note that in most tissue systems there are many cells interacting at once, and that there are likely multi-cell feedback loops in play which confound easy interpretation of such patterns.

To accommodate this data architecture and identify strongly perturbed edges regardless of category, we defined a ‘perturbation score’ that is calculatable for each edge, expressed as the product of the absolute values of the log-fold change for both the receptor and ligand, or:

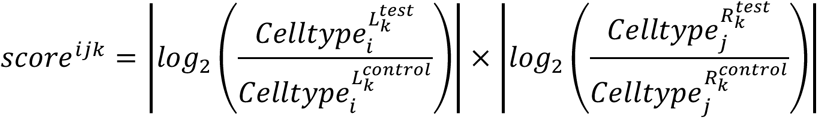

where 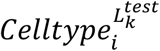 and 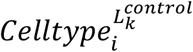 are the normalized expression of the ligand in celltype (i) in the test and control condition, respectively, and 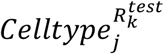 and 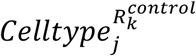 are the normalized expression of the receptor in celltype (j) in the test and control condition, respectively. The perturbation score is calculated in the above fashion so that edges with large changes on both the ligand and receptor side, regardless of sign, have very high scores, and minimally perturbed edges have a score of zero. Figure 3 demonstrates this concept in the interferon-simulated vs. control dataset of human peripheral blood mononuclear cells available through *SeuratData* (*13*). Figures 3C shows log fold changes for selected ligands and receptors, and the resulting perturbation scores (which are always positive) for each cell-ligand-receptor-cell edge. Perturbed edges can then be grouped by their differential pattern and their perturbation scores visualized in readily-explorable network form (Figure 3D).

### Longitudinal connectomics for a dynamic tissue-system process

Comparing tissue-level connectomics datasets can be difficult when the cell nodes are not directly comparable between the two tissue systems. Such a situation can occur if new cell types are recruited into, or eliminated from, a tissue system. Alternatively, normal differentiation processes may take place which cause a new cell type to emerge. Such changes occur frequently, whether due to inflammation, response to injury, wound healing, or normal development. Many unanswered questions in tissue science center around how these changing cellular landscapes correlate with shifting cell-cell communication patterns that are present in tissues and organs. Because the centrality analysis shown in Figure 2 quantifies network topology while being agnostic to specific nodes being present, this same technique allows for the comparison of disparate tissue systems which do not necessarily contain the same cell types.

As an example of a comparison of tissue systems containing different cell types, the muscle wound-healing dataset recently published by De Micheli et al. (*14*) provides an excellent use-case for longitudinal tissue connectomics. In this study, done *in vivo* in mice, muscle tissue was injured and then allowed to heal over time. scRNAseq was performed on Day 0, immediately before injury, and then on Days 2, 5, and 7 post-injury. Cells were parcellated on a per-time-point basis, leading to clear trends in cell type tissue fractions over time (Figure 4A). Certain cell types are present in the tissue throughout the wound healing process (i.e., endothelial cells, fibro/adipogenic progenitors (FAPs)), while others are recruited into the tissue solely during acute wound healing (i.e., monocytes/macrophages/platelets).

**Figure 4:**
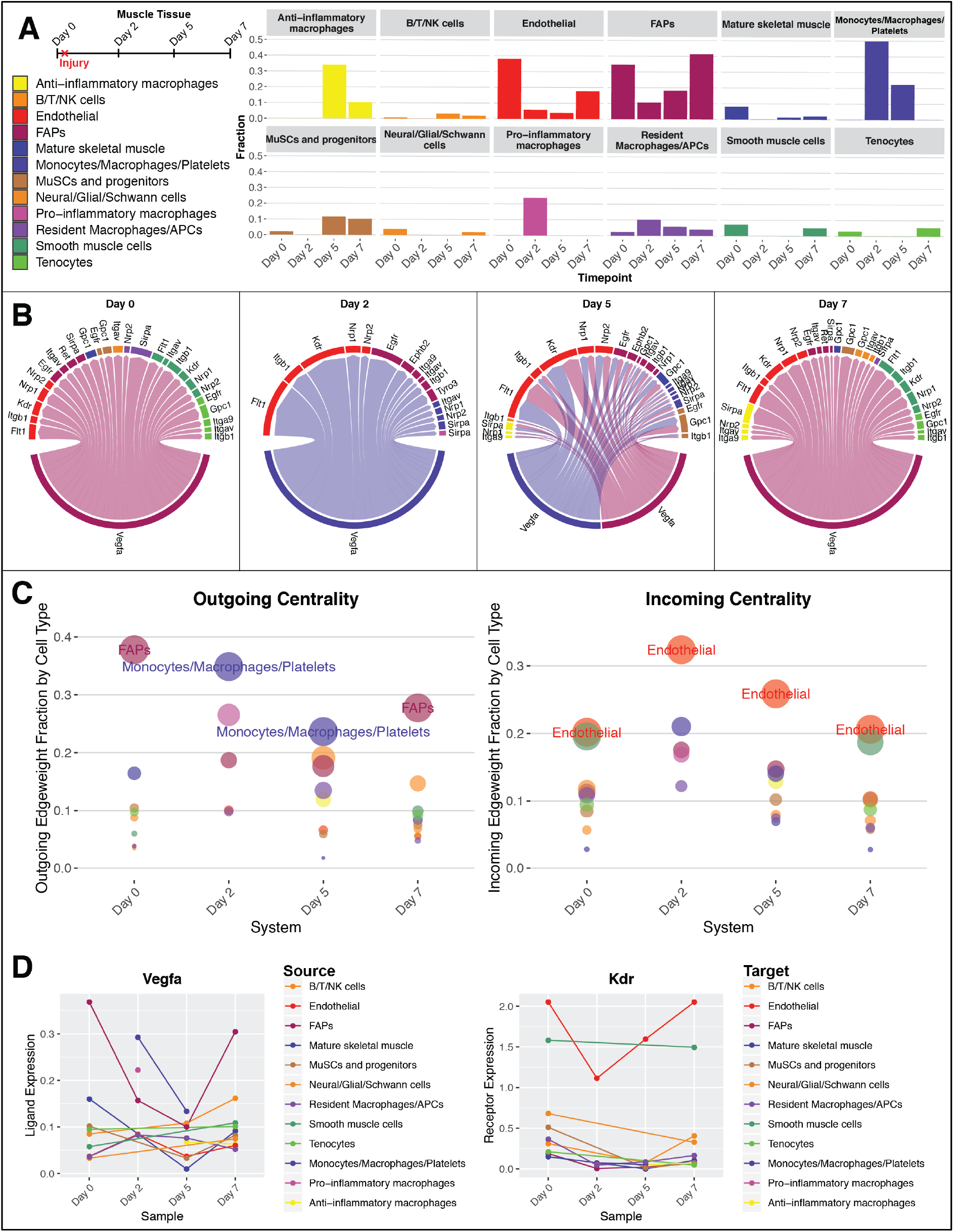
VEGFA signaling over time in healing muscle tissue. (A) Schematic of *in vivo* muscle injury experiment from De Michelis et al. (*14*) and associated dynamics in cell type dissociation fraction. (B) Vegfa signaling networks within muscle tissue at each time point. Edges coming from fibro/adipogenic progenitors (FAPs) are colored maroon while edges coming from monocytes/macrophages are colored blue. These network plots have been thresholded (minimum 10% expression) for legibility: the only source nodes which meet this criteria are the FAPs in Day 0, 5, and 7 and the Monocyte/Macrophage cluster in Days 2 and 5. Edge thickness is proportional to weight_scale_. (C) Centrality analysis, over time, for all Vegfa-mediated edges between all cell types, without any thresholding. Recruited monocytes take over dominant Vegfa production in healing tissue from homeostatic fibro/adipogenic progenitors. Endothelial cells are consistently in the dominant position to receive information, through their expression of multiple receptors including Kdr, Flt1, and Nrp1/2. (D) Normalized expression of Vegfa (ligand) and Kdr (receptor) for all cell types over time, showing relative expression. This analysis can be generalized to any network made from any combination of ligand-receptor mechanisms. For simplicity we show an example here based on only a single ligand.

For demonstration, we explore the network topology of cell-cell signaling based on Vascular Endothelial Growth Factor A (Vegfa) over time. Figures 4B-D suggests that there is a dramatic change in the dominant source of this ligand during wound healing. In the baseline state, FAPS are the dominant producers of Vegfa. Immediately after injury, FAPs reduce their production of this ligand, and newly recruited Monocytes/Macrophages/Platelets take up dominant production of Vegfa. On Day 5, these two cell types share this functional role, and at the conclusion of healing, FAPs again dominate the network. Endothelial cells, meanwhile, are always expressing a panoply of receptors for this secreted factor and are in a prime position to receive angiogenic information. We see that supporting cell types, including smooth muscle cells and tenocytes, are also in a position to receive Vegfa-mediated information before injury and after the conclusion of healing (green sectors). Further, we observe that muscle stem cells (MuSCs) are also capable of sensing aspects of Vegfa-mediated signaling, in particular on Day 5 post-injury, when they co-express and upregulate established angiogenesis-modulating receptors Egfr, Gpc1, and Itgb1 (*15–17*). It should be noted that, because of the parcellations chosen, this technique cannot necessarily tell the difference between an entire population of cells shifting in character versus a new phenotypic archetype emerging within an existing population. Further sub-clustering (i.e. finer, follow-up parcellation) is currently required to explore these kinds of questions.

### Comparison to existing cell-cell interaction software

There are multiple approaches to performing connectomic ligand-receptor mappings in single-cell data (*5–8*). In order to demonstrate that the mappings performed by *Connectome* are not spurious and that the edgeweights align with alternative methods of interaction weighting, we performed a cross-comparison with the software package *CellPhoneDB* (*6*). Selected results from this study are shown in Figure S3; the methods and workflow for this cross-workflow comparison are present in the Methods and in Supplemental Data D1, respectively. *CellPhoneDB* uses two mathematic techniques to assess interaction relevance, the first being a mean of the normalized values of ligand and receptor expression, and the second being a system-wide permutation test performed on the network for each ligand-receptor mechanism (*6*). These methods contrast notably with the *Connectome* edgeweights described above. Nevertheless, the two software packages yield highly comparable results and edgeweight rankings, both in terms of which mechanisms are most highly weighted for a given celltype-celltype pair, and in terms of which celltype pairs most preferentially engage through a given mechanism. The observed similarity in system architecture seen in Figure S3 both validates the relevance of the edgeweights calculated by *Connectome* and demonstrates that each technique converges on a common portrait of cell-cell signaling present in the underlying single-cell data.

## Discussion

*Connectome* is a multi-purpose toolset which can be used to map, explore, and visualize patterns of ligand-receptor expression in any single-cell dataset. The software is available open-access on GitHub and includes vignettes to replicate the full analysis presented here. It allows quantification and observation of both fine-grain (single-mechanism, single-celltype-to-celltype vector), and coarse-grain (total signaling family, cross-system) connectivity patterns.

There are a number of clear caveats to the above described techniques. First and foremost, strongly paired ligand and receptor expression is not direct evidence of cell-cell communication. For many ligand-receptor mechanisms, cells must be in direct or very close proximity in order to communicate via that mechanism. Suspension-based single-cell sequencing, however, does not preserve spatial histologic information, and so the techniques described here are attempting to make a somewhat qualitative portrait of a tissue based on what interactions are *possible*; any claim of actual transduction requires extensive wet-lab experimentation. Even for two cell types in a system which exclusively express a ligand and the cognate receptor each, respectively, a robust claim of transduction cannot be made: such an argument is only supported by evidence of downstream, intracellular effect via the receptor in question. Although this can be computationally predicted (*5*), the complexity of intracellular signaling networks makes this task fundamentally confounded for many ligand-receptor mechanisms.

Secondly, the outputs of *Connectome*, like any ligand-receptor mapping software, are only as good as the ground-truth database against which the original mapping takes place. This is why we have made the ground-truth customizable in this platform. Customization allows removal of extraneous connections with low biologic relevance, or addition of newly-discovered or researcher-determined mechanisms. The software here utilizes the FANTOM5 database without modification, but iterative application has shown that a custom database is generally required for many specific researcher inquiries. For immune cell studies in particular, it can be useful to curate a custom ground-truth database, which includes all immunomodulatory cues of potential interest.

*Connectome* is designed to be a fundamental tool for single cell researchers, computational biologists, and tissue engineers. It is intended to allow rapid, low-computationally-intensive-access to cell-cell signaling patterns that are present in single-cell data. Our intent is to allow researchers to quickly identify strongly-expressed signaling genes, to find strong pairings between cell types within identified signaling mechanisms, and to condense large amounts of network-level connectivity information into simple, quantitative plots which reflect the structure of tissue systems. We show here that *Connectome* can be applied to individual tissues, paired experimental conditions, and longitudinal datasets. In each case*, Connectome* yields biologically relevant information that can be used to help answer, and inform, specific questions regarding biological systems. We hope that it will prove useful for the larger biologic, engineering, and medical community.

## Acknowledgements

This work was supported by NIH grants F30HL143906 (Raredon), F30HG011193 (Garritano), 1R01HL138540 (Niklason), U01HL145567 (Kaminski), U54CA209992 (Levchenko), T32GM007205 (to M.S.B.R., J.G., and K.L.L.), R01GM131642, P50CA121974 and UM1DA051410 (Kluger) and DoD grant W81XWH-19-1-0131 (Schupp).

The authors would like to acknowledge Yifan Yuan for helpful conversations throughout software-craft and drafting and the Yale MSTP Office for their support throughout this collaboration.

## Author Contributions

Conceptualization, M.S.B.R, J.S., T.S.A., N.K., A.L., Y.K. and L.E.N; Methodology, M.S.B.R, J.G., J.Y., D.K., J.S., and Y.K.; Software, M.S.B.R., J.G., and J.Y.; Validation, M.Y. and J.Y.; Investigation M.S.B.R., J.Y., A.G., and K.L.L.; Writing – Original Draft, M.S.B.R.; Writing – Review & Editing, M.S.B.R., Y.K., N.K., A.L., and L.E.N; Supervision, Y.K., N.K, A.L., and L.E.N; Funding Acquisition M.S.B.R., J.G., J.S., N.K., A.L., Y.K. and L.E.N.

## Supplemental Methods

### Definitions

This manuscript uses some terms which are worth defining here. The term *connectome* refers to the complete set of interactions between nodes in a cell system, in the same way that *transcriptome* refers to the complete set of transcribed genes within a cell. The term *parcellation* refers to the way that the above system is divided up into distinct nodes, and in the application described in this manuscript, is synonymous with *celltype cluster*. The way a system is parcellated strongly affects its node architecture and therefore the shape of the resulting connectome. An *edge*, here, is a single unique celltype-ligand-receptor-celltype interaction. *Edge attributes* are quantitative or qualitative pieces of information associated with an edge. A *differential connectome* is the network that results when two connectomes are directly compared, edge-for-edge. *Centrality*, as used in the text, refers to quantitative metrics of how ‘connected’ a given node is to other nodes, in either an outgoing (sending) or incoming (receiving) fashion.

### Edgeweight definition and usage

The *Connectome* software automatically refines two edgeweights which are meant to be employed in distinct use-cases and are each useful for specific computational explorations and illustrations. The ‘weight_norm’ edge attribute is derived from the normalized expression of the ligand and the receptor in the single-cell data. This edge weight is meant to be used when comparing multiple distinct cellular systems to each other, as when computing differential connectomics. The ‘weight_scale’ edge attribute is derived from the z-scores of the ligand and the receptor in each edge, and is of higher value when the ligand and receptor are more specific to a given pair of cell types. This edgeweight is meant to be used when exploring a single cell system, but it is not ideal for direct edge-to-edge cross-system comparisons, since variable cell sampling and/or cell parcellation will affect it.

It should be noted that batch effects have the potential to affect normalized gene expression values (*18*), and therefore also have the potential to affect certain quantitative edge attributes in the connectomic mapping. Care should therefore be taken when comparing connectomics between systems with substantial batch effects.

### Calculation of statistical significance (single tissue system)

For certain computational and visualization tasks, it becomes advisable to narrow focus from the full connectome to only those edges deemed significant. To do so, we recommend limiting analysis to only those edges where the ligand and the receptor are both expressed in greater than a certain fraction of their respective clusters, generally 10%. It is often then further useful to limit analysis to those edges where both the ligand and the receptor have a p-value of less than 0.05, as determined by a system-wide Wilcoxon Rank Sum Test (calculated by default within *CreateConnectome*.) These thresholds can be further reduced within *FilterConnectome,* to identify those pathways of greatest significance, and can be crossed with additional thresholding to limit edge representation to specific vectors or signaling families (see vignettes online).

### Calculation of statistical significance (differential tissue system)

In the case of differential connectomics, special care has to be taken to include edges which are not significant in a single experimental condition but which become significant in another. The min.pct argument in *DifferentialConnectome*, therefore, only needs to be satisfied in either the reference dataset or in the test dataset; this allows inclusion of mechanisms which emerge or disappear.

The vignette online shows how to use a Wilcoxon Rank Sum test to compare each pair of commensurate cells across two datasets and to thereby determine which ligands and which receptors are differentially expressed to a statistically significant degree. The differential connectome can then be thresholded to only those edges where both the ligand on the sending cell and the receptor on the receiving cell are differentially expressed between experimental conditions. All differential connectomics presented and discussed in this manuscript and online utilize this statistical approach.

### Centrality definition and usage

*Centrality* and *CompareCentrality* both calculate and plot-to-compare two key centrality metrics for networks of interest: the cumulative incoming/outgoing edgeweight for each node, and the Kleinberg Hub and Authority scores for each node. These two centrality metrics often correlate, but they do not always flawlessly agree on exact node ranking. There is no single, proper and agreed upon way to best calculate network centrality (*19*); these two metrics were chosen because they are commonly used in network research and are biologically interpretable. Cumulative incoming/outgoing edgeweight fraction is the sum of the incoming and outgoing edges, calculated per node, expressed by default as the fraction of total edgeweight within the considered system; this ranks each node’s ability to contribute or listen to a given set of interactions. Hub-Authority centrality is a heavily-used information flow metric (*20*), which should be interpreted, in this instance, to reveal Hubs which send signal to Authorities, and Authorities which receive information from Hubs. Each of these metrics provide interpretable information regarding biologic network structure.

### Perturbation score definition and usage

The perturbation score, defined explicitly in the main text, varies from 0 to Infinity, and increases proportionally to both the log fold-change of the ligand (in either direction) and to the log fold-change of the receptor (in either direction). It is generally useful to limit analysis to those edges which have ligand and receptor expression in > 10% of their respective clusters in either the control or the test condition, to avoid noisy measurements (see Supplemental Data D1); this thresholding functionality is built into *CircosDiff* and *DifferentialScoringPlot*.

### Signaling family categorization

The signaling family categorizations used in this manuscript are carried over from (*1*) with minor updates per recent literature findings. These groupings are loaded by default within *Connectome.* It should be noted that each signaling mechanism is allowed only a single designation in this formulation; in biologic reality, however, many signaling mechanisms can be considered to belong to multiple signaling families. These groupings are meant as a guide for hypothesis generation and later downstream exploration, rather than as definitive classification.

### Analysis of pancreas data

A fully replicable workflow for the processing of the pancreas data and generation of subpanels in Figure 1 and Figure 2 is included in Supplemental Data D1. In brief, the panc8 data was downloaded from SeuratData, normalized, scaled, and run through the *CreateConnectome* with a min.cells.per.ident cutoff of 75. Downstream analysis was limited to those edges where both the ligand and the receptor were expressed in > 10% of their respective clusters, and which had a p-value < 0.05 for both the ligand and the receptor. Additional thresholds applied for specific visualization purposes present in the figures can be seen in Supplemental Data D1.

For the centrality analysis present in Figure 2, *Centrality* was first run across all signaling families. Each signaling family was then grouped according to which of the 4 cell classes was found to dominate incoming centrality within the given dataset. *Centrality* was then re-run 4 times, once for each set of signaling families, to yield the 4 sub-graphs. A script to replicate this workflow is present in Supplemental Data D1.

### Analysis of IFN-stimulated vs. Control PBMC data

A fully replicable workflow for the processing of the IFN-stimulated vs. Control PBMC data and generation of subpanels in Figure 3 is included in Supplemental Data D1. In brief, the ‘ifnb’ dataset was loaded from *SeuratData.* Each dataset was normalized, scaled, and run through *CreateConnectome*. The two connectomes were then passed to *DifferentialConnectome.* A Wilcoxon Rank Sum test was then performed for each individual cell across conditions, for all ligands and all receptors, and the results from this test were used to limit the differential connectome to only those edges showing statistically significant differences (p < 0.05) for both the ligand and the receptor. Downstream analysis was further limited to those edges which had ligand and receptor expression in > 10% of their respective clusters in *either* the control or the test condition. Figure 3C was made with *DifferentialScoringPlot* and Figure 3D was made with *CircosDiff*, both of which are built in to *Connectome.*

### Analysis of muscle wound-healing data

A fully replicable workflow for the processing of the muscle wound-healing data and generation of subpanels in Figure 4 is included in Supplemental Data D1. In brief, raw data was loaded from De Micheli et al. (*14*), normalized, scaled, and run through *CreateConnectome*. *ggplot* was use for Fig 4A, *CircosPlot* for Fig 4B, and *CompareCentrality* for Fig 4C. A custom script, present in Supplemental Data D1, was used for Fig 4D.

### Comparison between *Connectome* and *CellPhoneDB*

To create Supplemental Figure S3, the most recent version of CellPhoneDB (v2.1.4) was downloaded and the log normalized pancreas gene expression data and corresponding cell type labels were input to CellPhoneDB. All the CellPhoneDB parameters were set to default. Edge were only considered which had their mechanisms present in the FANTOM5 database, their ligand and receptor both expressed in greater than 10% of their respective clusters, and a p-value < 0.05 (in the case of *CellPhoneDB*) or a ligand and receptor p-value of < 0.05 (in the case of *Connectome).* For visualization, those mechanisms shown in Figure S3 are a selection of those, within the above set, which also have a ligand and receptor z-score > 1. The function *EdgeDotPlot*, provided within the *Connectome* software package, was used to make panel A.

## Supplemental Figures

**Figure S1:**
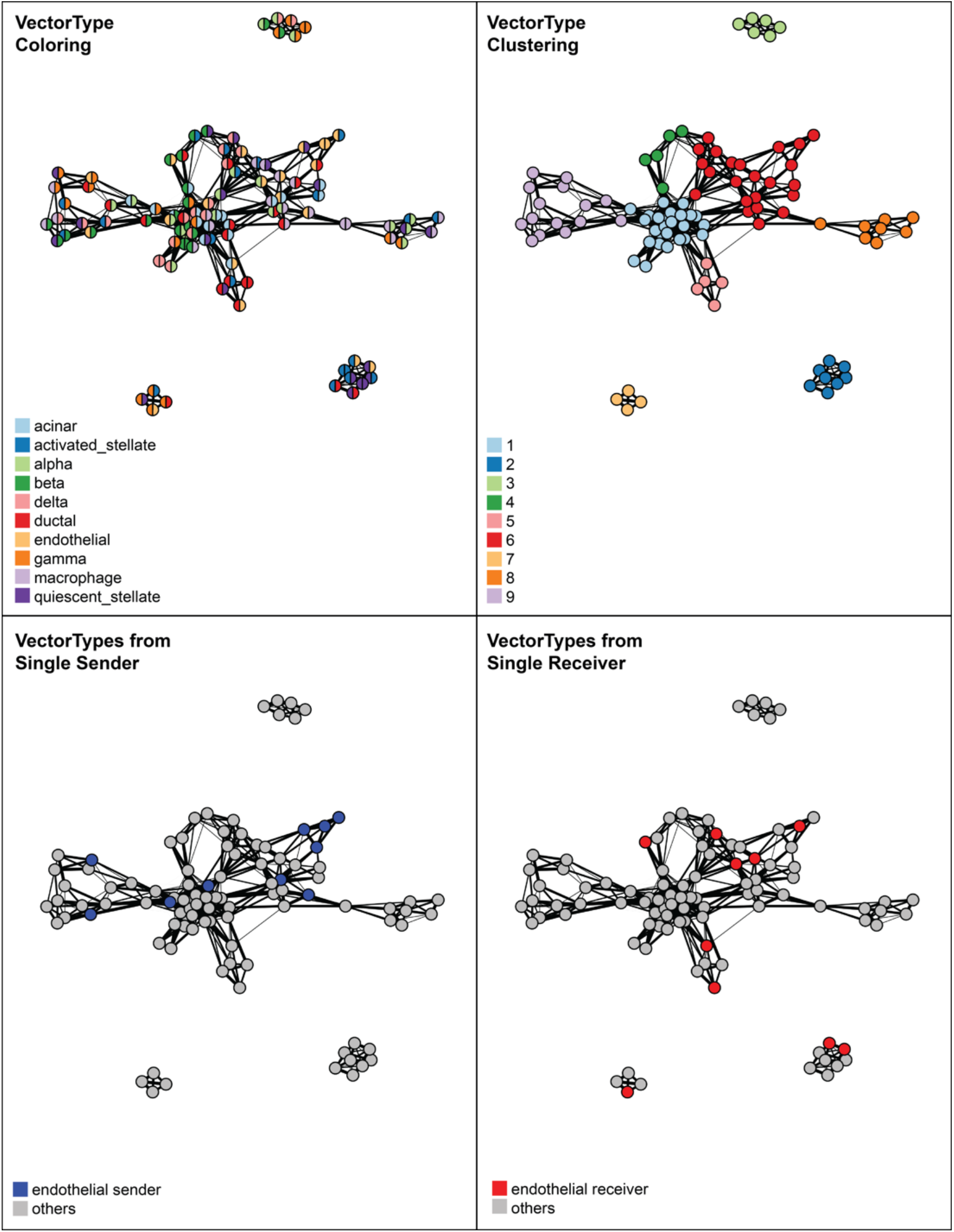
Shared nearest neighbor graph visualization of vector types. Graphs shown are generated using k=3 in k-nearest neighbor calculation which uses the weight_norm edge attribute of all ligand-receptor mechanisms as features. Each node is a vector type, and an edge is drawn between two nodes that share at least one neighbor. The weights of the edges that reflect the similarity between two nodes are calculated using the rank scheme defined by (*21*). Nodes in the upper left graph are colored in two parts, with the left color representing the sending celltype and the right color representing the receiving. The upper right shows a Louvain clustering of this same graph. As a demonstration, the lower panels emphasize those vectortypes originating only from endothelial cells and landing only on endothelial cells, respectively. Note (i.e. in Cluster 2) that two vectortypes can be highly similar even though they do not all originate from, or land on, the same cell types. This is indicative of substantial overlap in the ligand-receptor mechanism weighting between those two cell types.

**Figure S2:**
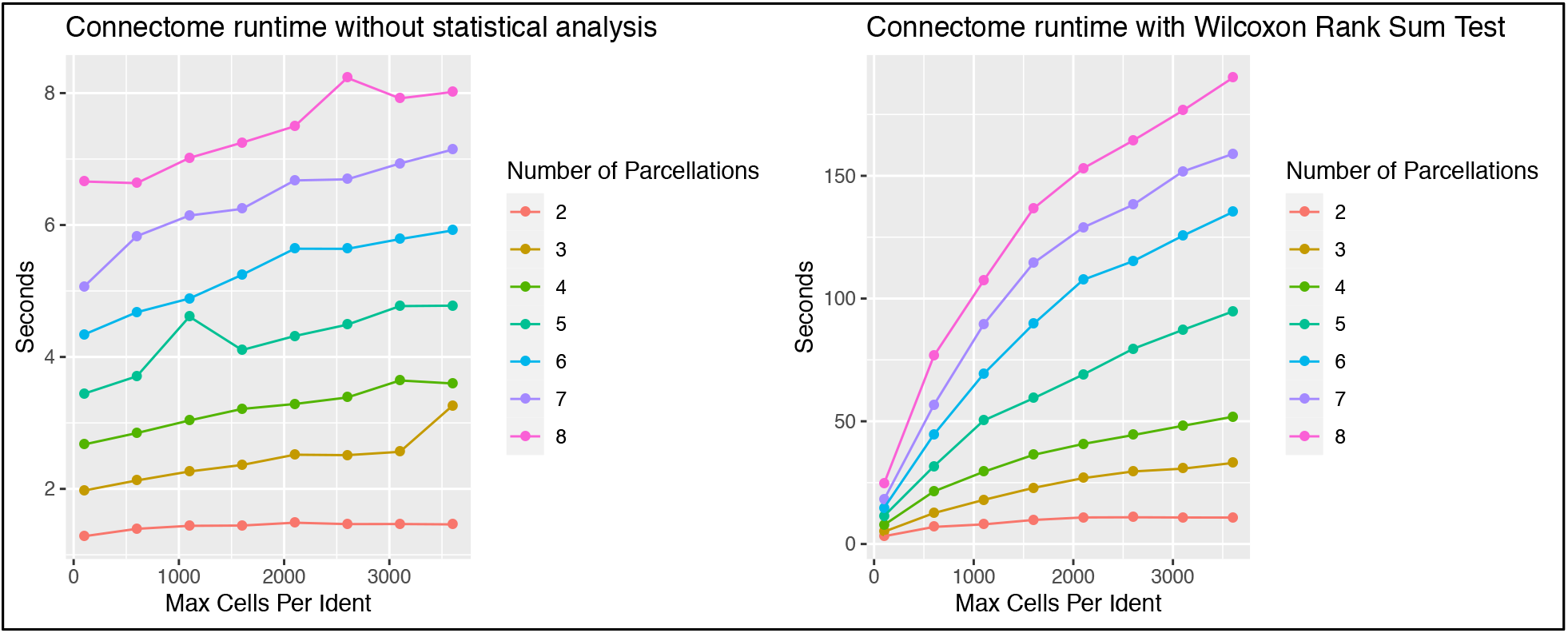
Runtime for connectome computation with and without statistical analysis. Simulation was run on the panc8 dataset from *SeuratData,* with and without calculation of p-values for each ligand and receptor, on each edge, via a system-wide Wilcoxon Rank Sum test. In both cases, runtime is dependent on both parcellation number and cell number. This simulation was run on a 2.9 GHz 8-core MacBook Pro laptop.

**Figure S3:**
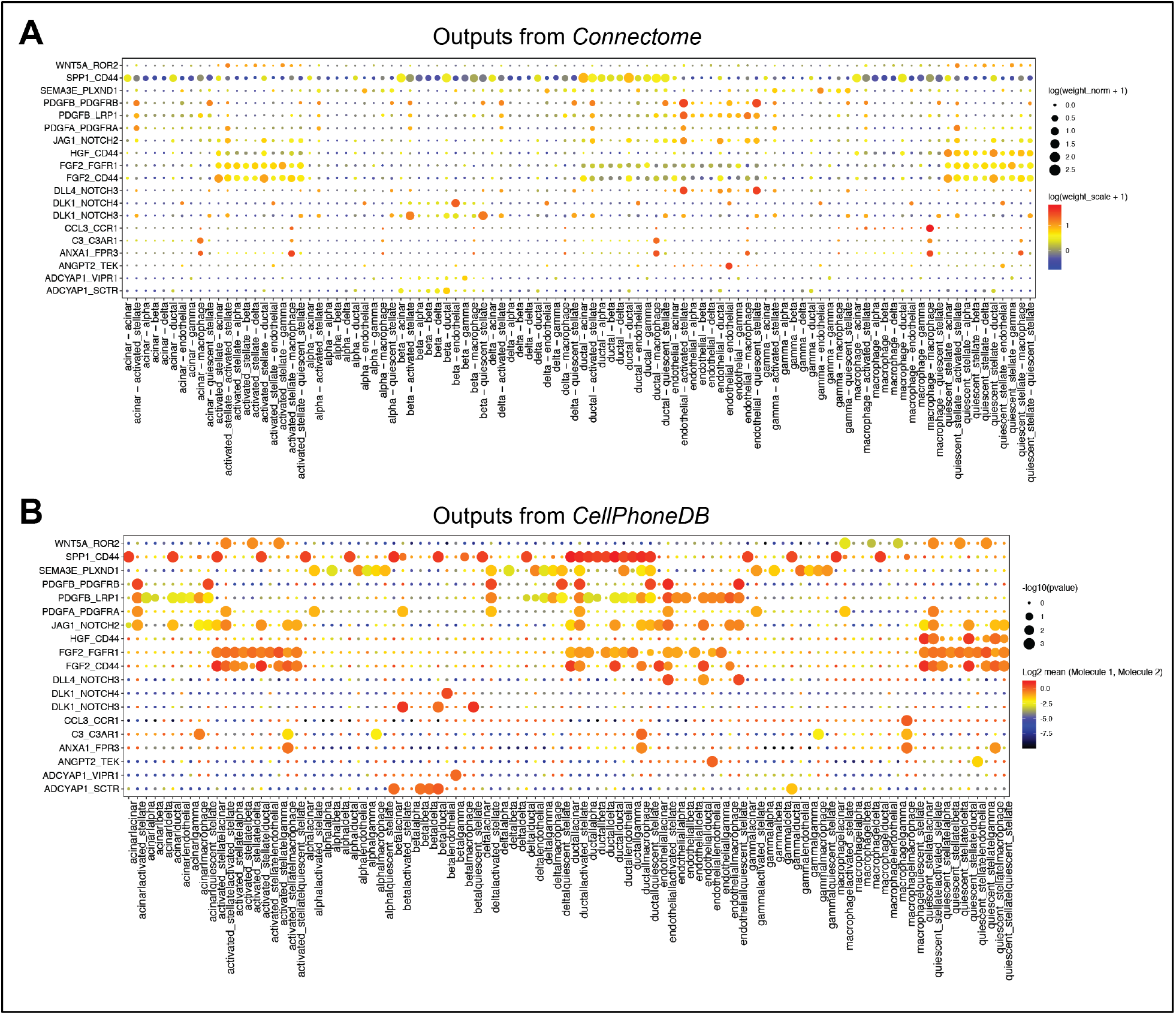
Comparison of outputs from *Connectome* with outputs from *CellPhoneDB.* To validate that the mappings and edgeweight calculations performed by *Connectome* are accurate, we compared A) selected outputs from the application of *Connectome* to the panc8 dataset to B) those generated through the application of *CellPhoneDB (6)* to the same dataset and with the same ground-truth list of ligand-receptor mechanisms (see Methods). Note the high similarity of findings across all columns and rows, in particular the specificity of DLK1-NOTCH4 to the beta – endothelial vector, the specificity of ANGPT2-TEK to the endothelial – endothelial vector, and the specificity of DLL4-NOTCH3 to the two endothelial – stellate cell vectors. The two plots are not identical because the two software packages use very different means of calculating cell-cell interaction relevance; we have attempted to make nonetheless comparable visualizations here. In (A), the size of the dot represents the log of the product of the normalized expression of ligand and receptor plus 1, while the color represents the log of the mean of the scaled expression of ligand and receptor plus 1. In (B), the size of the dot represents the negative log of the permutation-test p-value of each mechanism, while the color represents the log of the mean of the normalized expression of ligand and receptor. Note that the color of the dots in panel A (*Connectome* weight_scale_) closely correlates with the size of the dots in panel B (*CellPhoneDB* p-values). These plots are intended to convey that the two mappings and statistical weightings are similar and that both approaches reveal similar biological patterns.

## Supplemental Data D1: R scripts to replicate all figures in manuscript

All software is available at https://github.com/msraredon/Connectome. Detailed vignettes and instructions for use are published at https://msraredon.github.io/Connectome/. For ease of reproduction, the exact scripts used to generate the analysis in this paper are copied below.

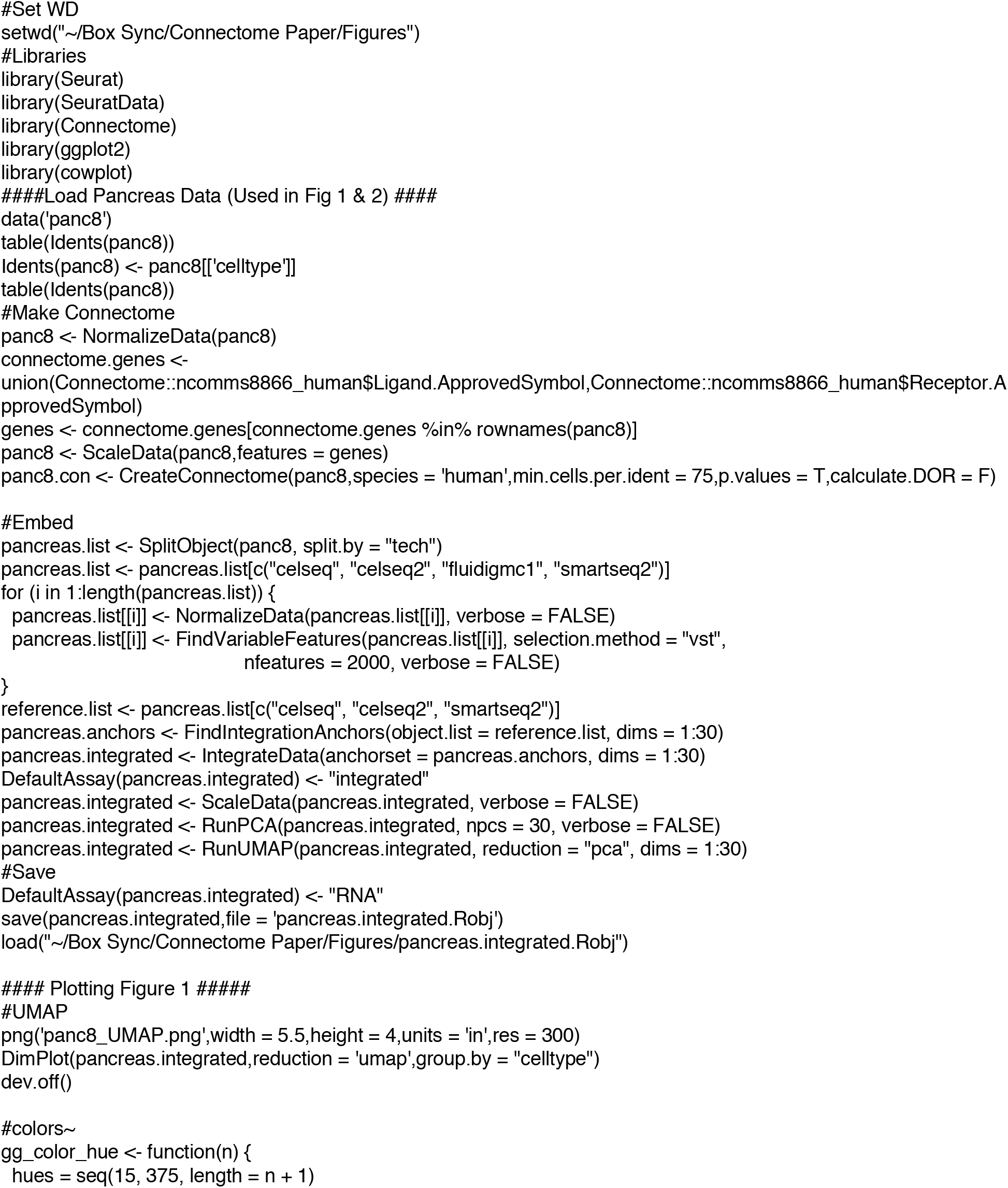
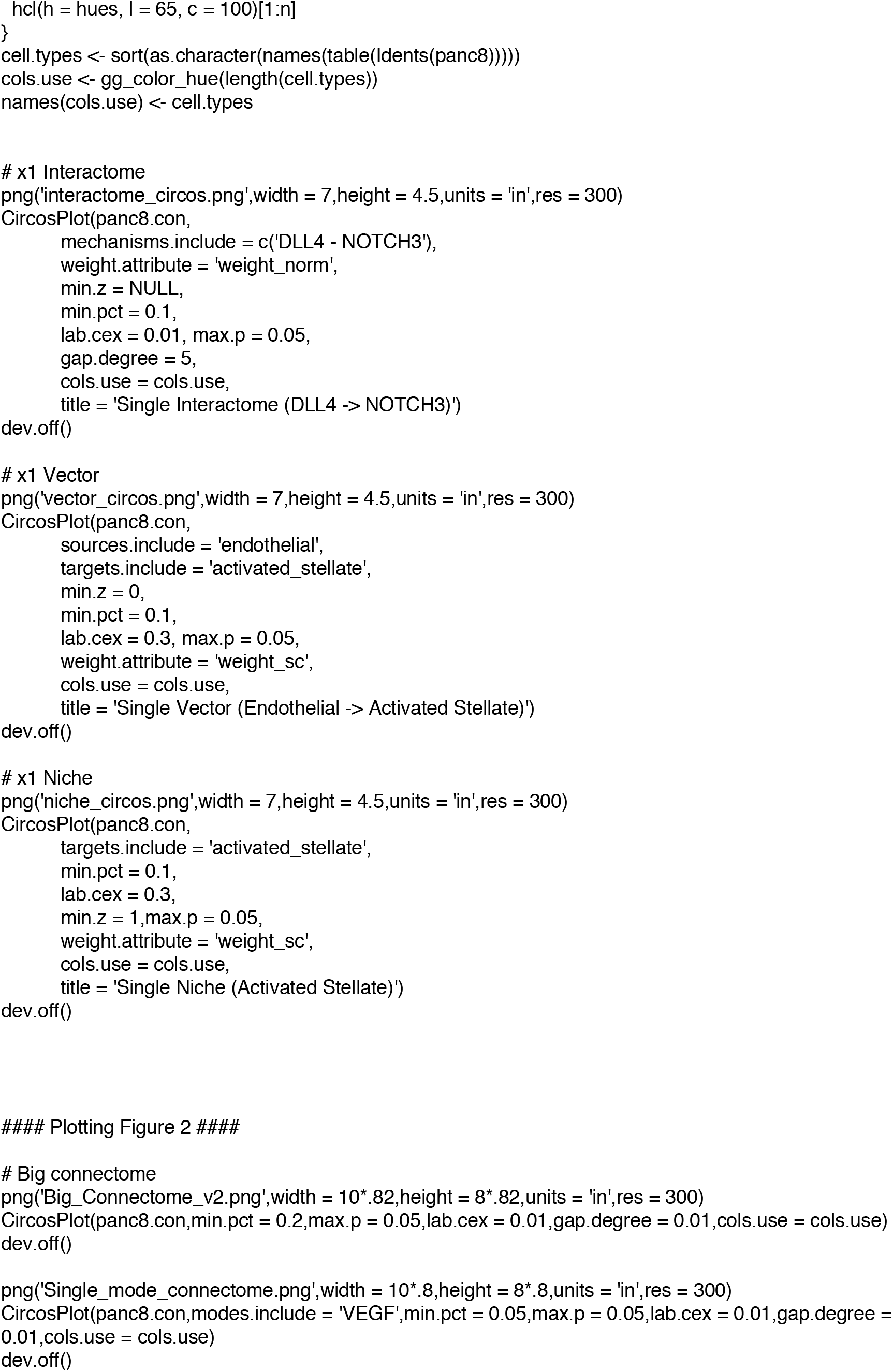
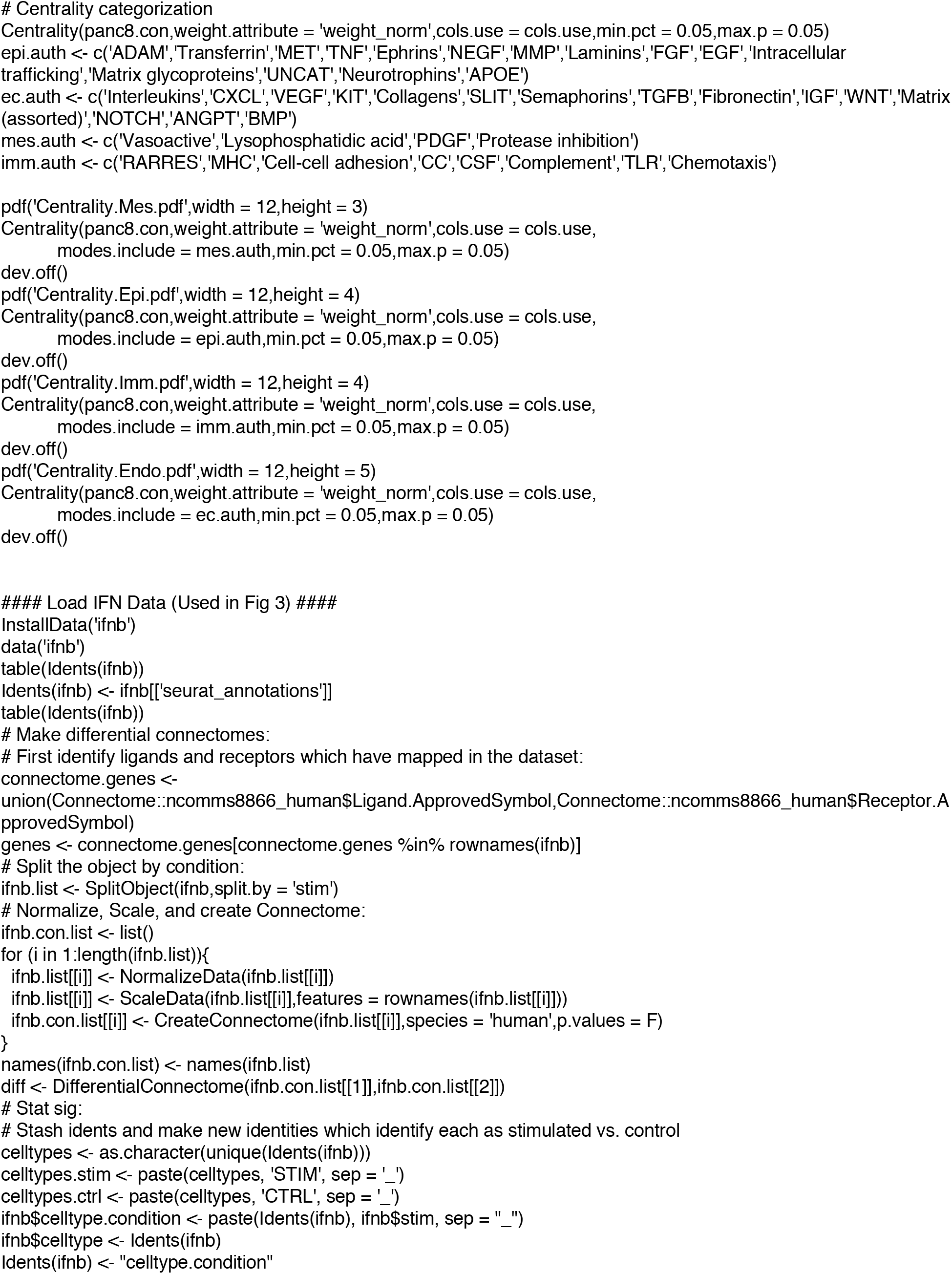
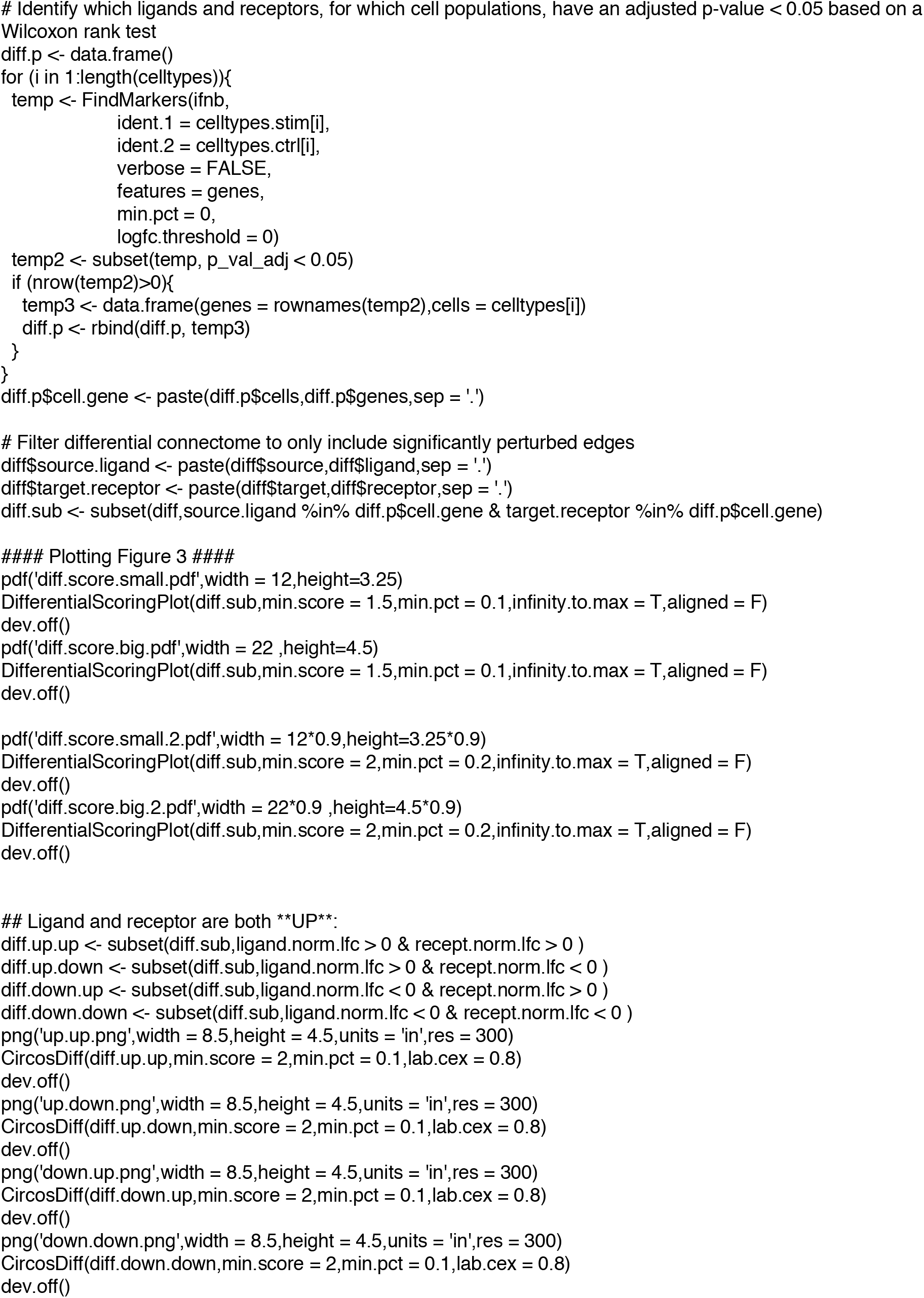
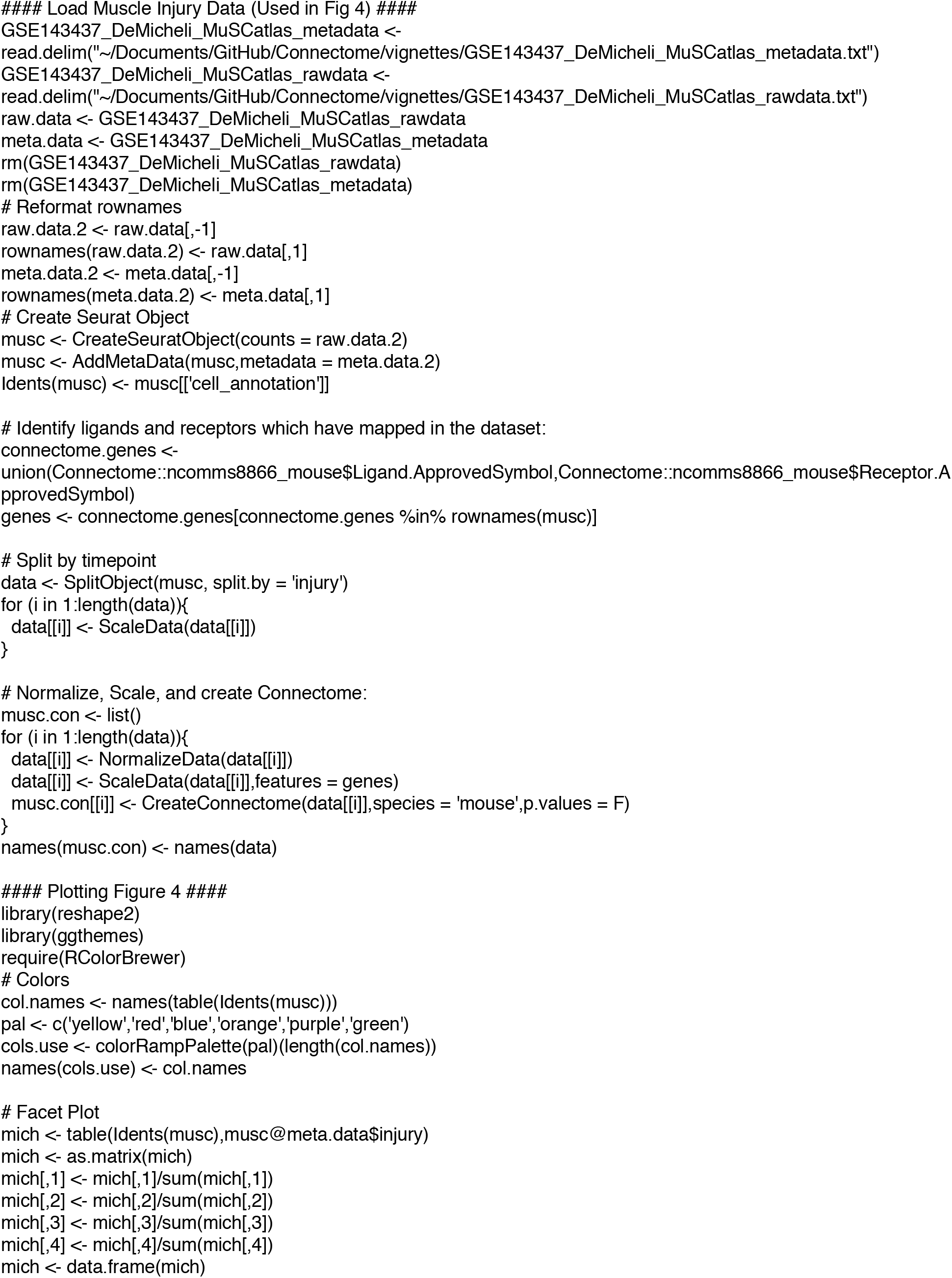
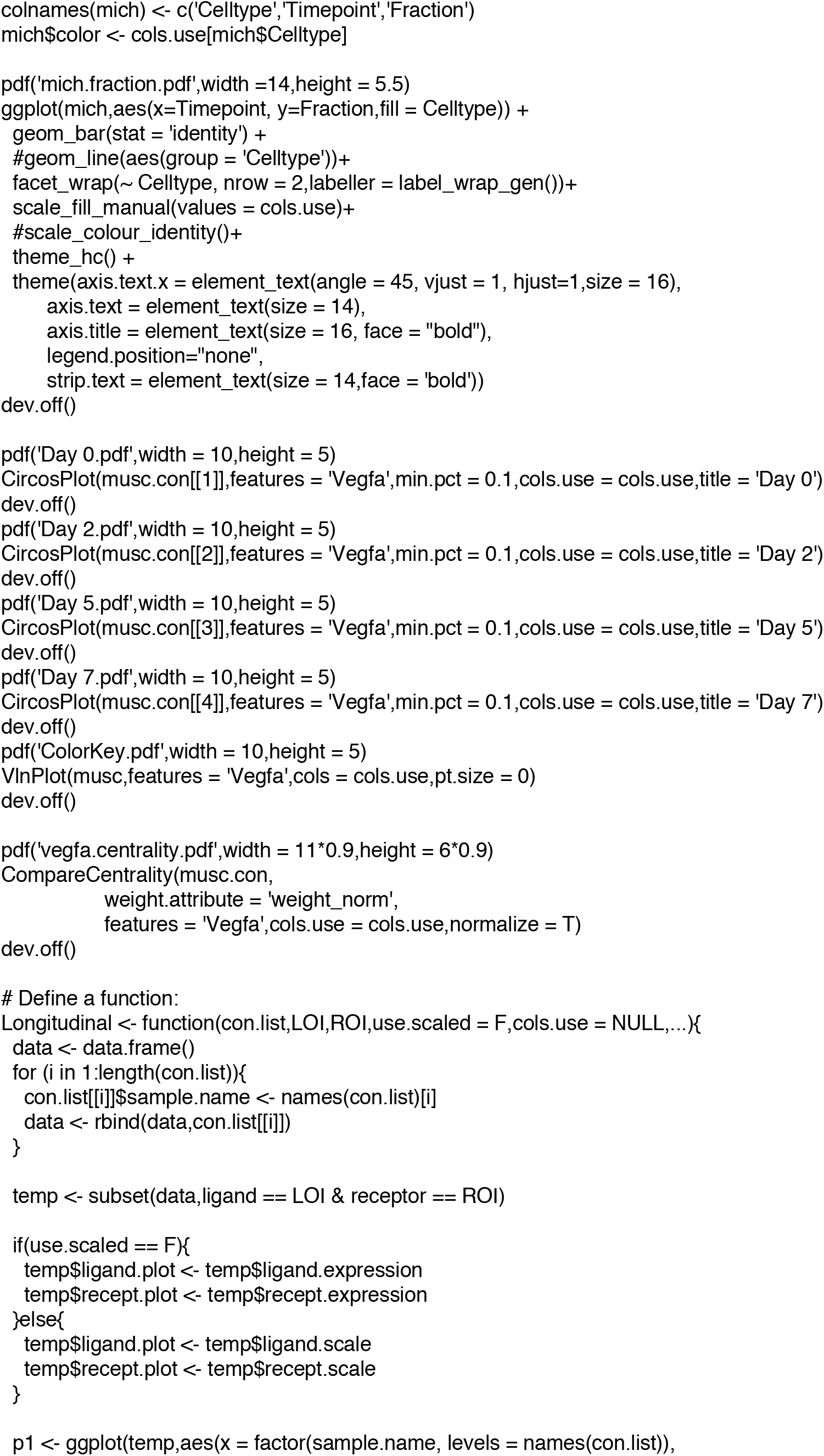
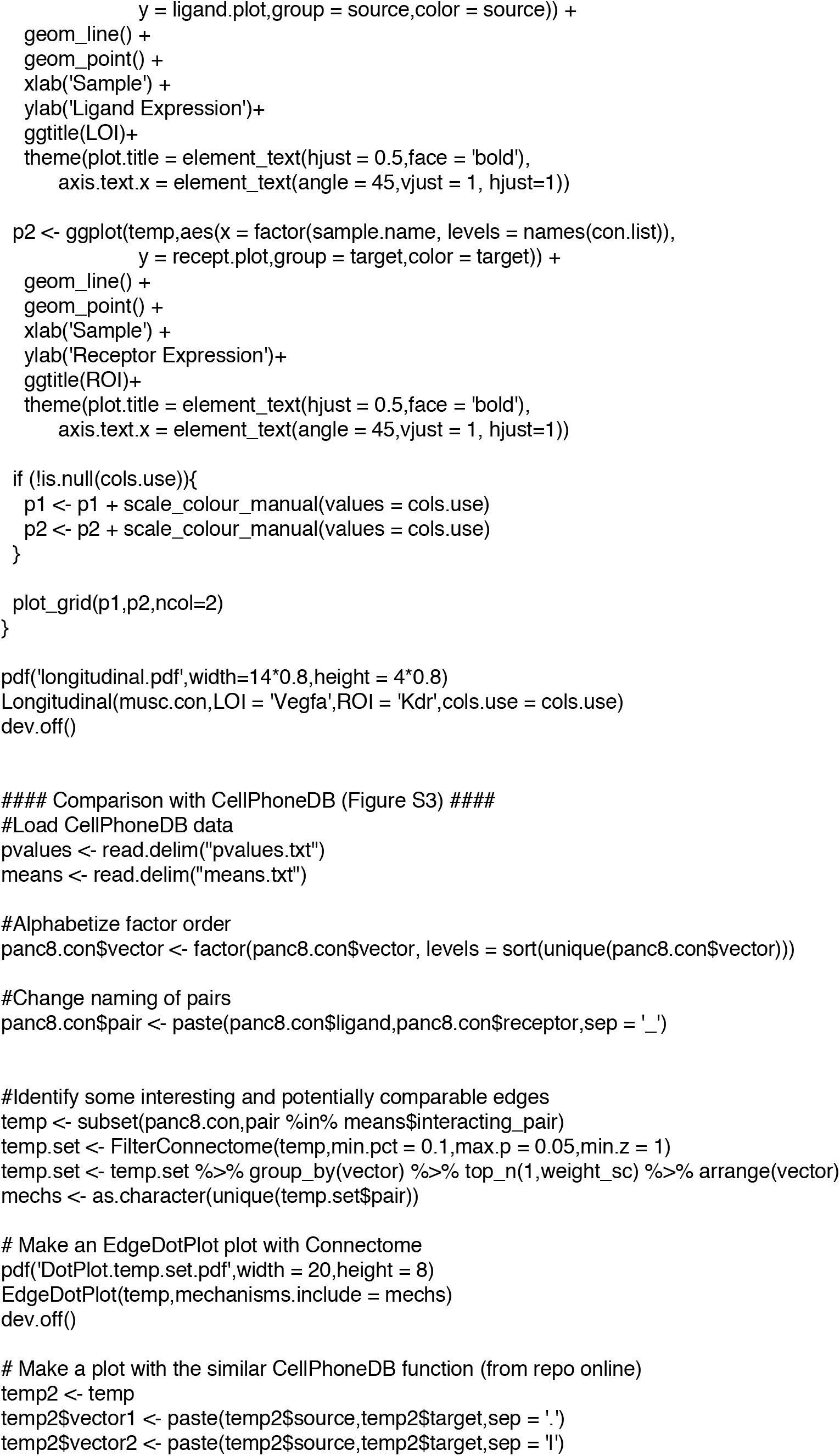
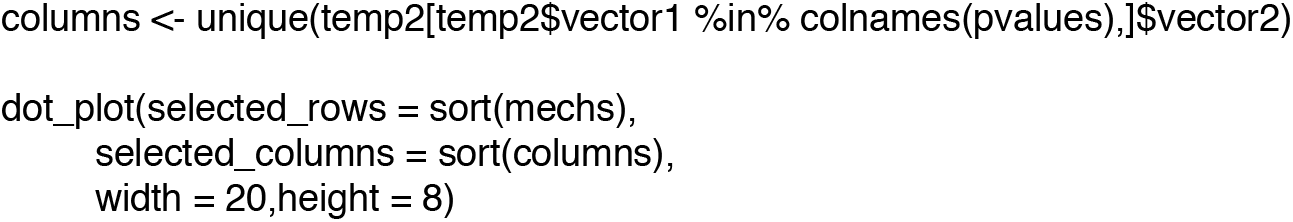

